# Cellular translational enhancer elements that recruit eukaryotic initiation factor 3

**DOI:** 10.1101/2021.07.15.452589

**Authors:** Jiří Koubek, Rachel O. Niederer, Andrei Stanciu, Colin Echeverría Aitken, Wendy V. Gilbert

## Abstract

Translation initiation is a highly regulated process which broadly affects eukaryotic gene expression. Eukaryotic initiation factor 3 (eIF3) is a central player in canonical and alternative pathways for ribosome recruitment. Here we have investigated how direct binding of eIF3 contributes to the large and regulated differences in protein output conferred by different 5′-untranslated regions (5′-UTRs) of cellular mRNAs. Using an unbiased high-throughput approach to determine the affinity of budding yeast eIF3 for native 5′-UTRs from 4,252 genes, we demonstrate that eIF3 binds specifically to a subset of 5′-UTRs that contain a short unstructured binding motif, AMAYAA. eIF3 binding mRNAs have higher ribosome density in growing cells and are preferentially translated under certain stress conditions, supporting the functional relevance of this interaction. Our results reveal a new class of translational enhancer and suggest a mechanism by which changes in core initiation factor activity enact mRNA-specific translation programs.

## Introduction

mRNA-specific translational activity – the number of protein molecules produced per mRNA – varies by orders of magnitude under normal growth conditions and is extensively regulated in response to a wide range of physiological signals (Ghazalpour et al., 2011; Lahtvee et al., 2017; Schwanhäusser et al., 2011; Vogel and Marcotte, 2012). 5′-untranslated regions (5′-UTRs) directly contact the translation initiation machinery and can strongly influence the rate of translation (Hinnebusch et al., 2016). For example, sequence differences between native yeast 5′-UTRs are sufficient to drive greater than hundred-fold differences in translation initiation (Rojas-Duran and Gilbert, 2012). Despite great progress towards illuminating the fundamental mechanisms by which eukaryotic mRNAs recruit ribosomes to initiate translation, many quantitatively large mRNA-specific differences in translation initiation remain unexplained.

For most cellular messages, translation initiation requires the concerted action of many eukaryotic initiation factors (eIFs). Cellular mRNAs begin with a 5′-m^7^G cap that is recognized by the eIF4E subunit of the cap binding complex, eIF4F. Ribosomes are recruited to mRNA as 43S pre-initiation complexes (PIC) that consist of a 40S small ribosomal subunit bound to eIF3, a ternary complex of eIF2•GTP•Met-tRNA_i_, and additional factors. During cap- and scanning-dependent initiation, the assembled PIC scans from 5′ to 3′ to find the start codon, at which point the 60S large ribosomal subunit joins and protein synthesis begins (Dever et al., 2016). Eukaryotic initiation factor 3 (eIF3) is a central player in this canonical pathway for translation initiation. eIF3 consists of five core subunits that are conserved from yeast to man with seven additional subunits present in filamentous fungi and multicellular eukaryotes (Cate, 2017; Valášek et al., 2017). Consistent with its large size and conservation, biochemical and structural studies reveal an extensive network of eIF3 interactions within the PIC, which stabilize the PIC and promote mRNA recruitment (Aitken et al., 2016; Valášek et al., 2017).

eIF3 is also required for several non-canonical or cap-independent modes of translation. In vitro, eIF3 enhances ribosome binding to model cellular mRNAs in the absence of a cap (Mitchell et al., 2010). Four of the conserved subunits, eIF3a, b, c and g, contain RNA-binding domains that may bind mRNA as well as 40S ribosomes during initiation (Sun et al., 2013). Consistent with this possibility, crosslinking and immunoprecipitation of eIF3 from human 293T cells identified hundreds of specific crosslink sites mostly within 5′-UTRs (Lee et al., 2015). Further characterization of one eIF3 direct target, c-JUN, showed that eIF3 bound to a structured 5′-UTR element and enhanced cap-dependent translation, although the majority of efficiently translated mRNAs did not crosslink to eIF3. Certain RNA viruses such as hepatitis C initiate translation without mRNA caps using structured 5′-UTR elements that bind to host eIF3 with high affinity (Filbin ME et al., 2009). Thus, direct binding of eIF3 to the 5′-UTR is sufficient to initiate downstream steps in translation initiation. Because this mode of ribosome recruitment is insensitive to regulatory mechanisms that target the activity of the cap binding complex, cap-independent and eIF3-dependent initiation is thought to play an important role in selective protein synthesis during times of stress (Gilbert, 2010; Shatsky et al., 2018). Whether high affinity binding to eIF3 is broadly significant for cellular translation activity was unknown.

Here, we have used an unbiased high-throughput approach to determine the affinity of the yeast eukaryotic initiation factor 3 towards 5′-UTRs from 4,252 genes and then compared direct eIF3 binding to translation activity in cells. We incubated purified yeast eIF3 with a synthetic pool of 5′-UTRs at a range of protein concentrations and sequenced the bound RNAs to identify hundreds of specific binders. Quantitative filter binding assays validated specific 5′-UTRs as high affinity binders. We identified a sequence motif, AMAYAA, that was significantly enriched within unstructured regions of 5′-UTRs that bound eIF3 and found that mRNAs containing this 5′-UTR motif show higher translation activity in rapidly growing yeast. mRNAs that bound eIF3 in vitro maintained higher translation activity under conditions of limiting eIF3 and also during glucose starvation. Together, these results suggest a broad role for direct eIF3 binding to specific 5′-UTR elements during normal and stress-responsive translation.

## Results

### A high-throughput assay for direct binding of eIF3 to specific 5′-UTRs

Because of the emerging role of eIF3 in mediating the translation of specific transcripts, we asked whether specific yeast mRNAs bind eIF3 with high affinity. Such direct binding to eIF3 could promote efficient translation in growing cells or contribute to selective protein synthesis during stress conditions where cap binding activity is downregulated. To survey the binding specificity of individual transcripts for eIF3 across the transcriptome, we performed RNA Bind-n-Seq (RBNS), a quantitative high-throughput assay for RNA binding affinity in vitro (Lambert et al., 2014). We designed an RNA library of sequences derived from deep sequencing of full-length yeast 5′-UTRs (Pelechano et al. 2014) (Methods). This pool included 5′-UTR sequences from 4,252 genes.

eIF3 was purified from yeast cells and its activity confirmed by biochemical complementation of translation extracts from temperature-sensitive *prt1-1* cells lacking functional eIF3 as previously described (Phan et al., 1998) (Figure S1a, b). Purified eIF3 at various concentrations (0-1500 nM) was incubated with the 5′-UTR pool and bound RNA:eIF3 complexes were separated from free RNA by passing through a nitrocellulose filter and used to prepare RNA sequencing libraries together with input RNA (Figure 1a). For each concentration, we determined an eIF3 enrichment score in the bound fraction as the relative frequency of a given RNA in the bound library compared to the input. The normalized reads for each eIF3 concentration were highly reproducible between replicates (Figure 1b and Figure S1c). The replicate scores were therefore averaged in all subsequent analyses. Averaged eIF3 enrichment scores from adjacent eIF3 concentrations were highly correlated above 55 nM eIF3 (Spearman R > 0.9, Figure 1c), which is an indicator of good quality RBNS libraries (Lambert et al., 2014).

**Figure 1:**
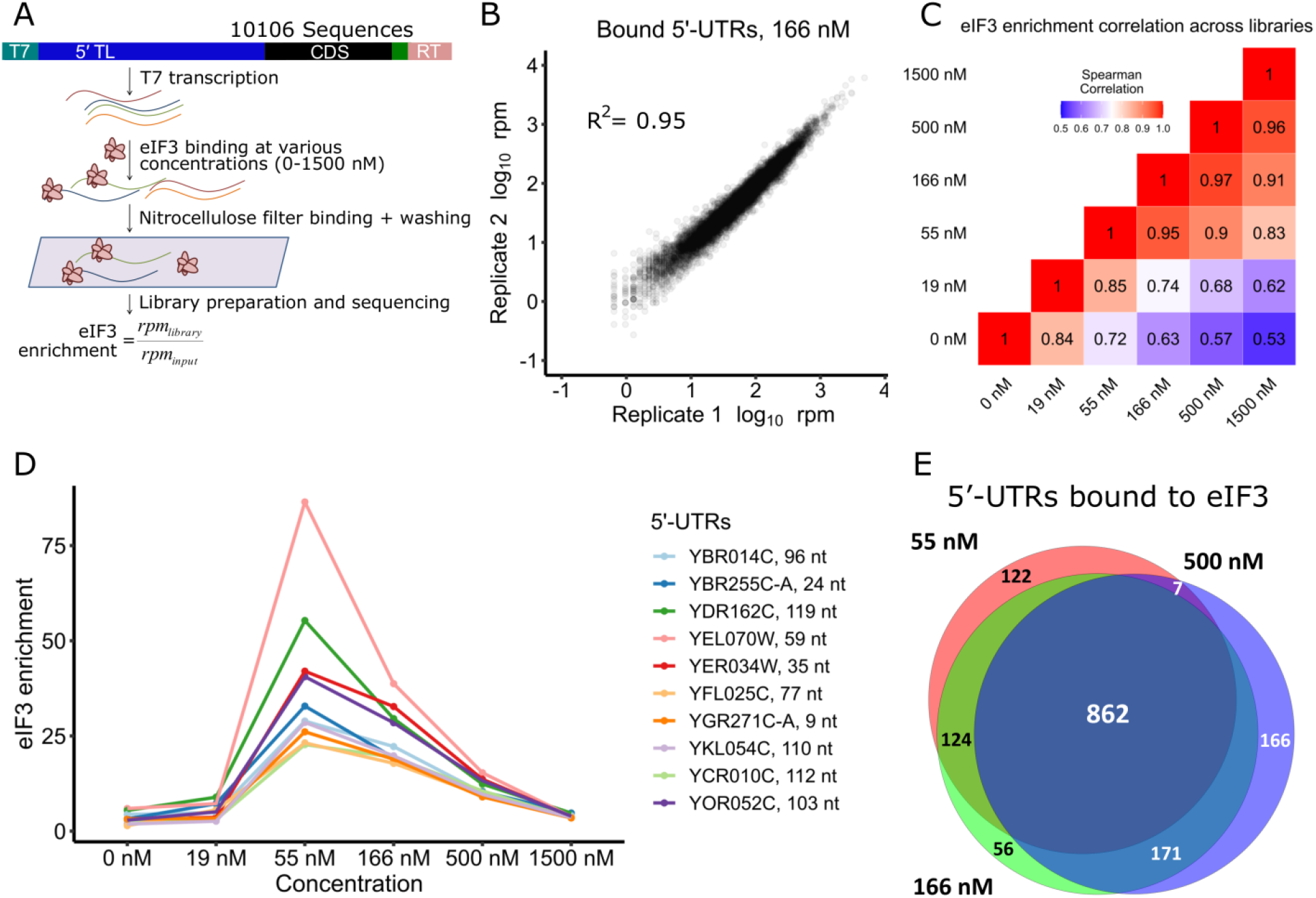
Hundreds of natural yeast 5′-UTRs bind to eIF3 specifically. **a)** Scheme of RNA Bind-n-Seq (RBNS) assay. Designed DNA oligos encoding ~10,000 yeast 5′-UTRs are transcribed in vitro. The resulting RNA pool is incubated with various concentrations of purified eIF3, and protein-bound RNAs are captured on nitrocellulose and sequenced. Enrichment is the frequency of a 5′-UTR in the bound sample normalized to input RNA. **b)** RBNS results are reproducible. Shown are bound reads in two replicates of 166 nM eIF3. **c)** Enrichment between 55 and 500 nM eIF3 was well correlated. **d)** Enrichment of the top 10 eIF3 binders from the 166 nM library traced across all libraries. Peak enrichment at 55 nM eIF3 is consistent with high affinity binding. Less enrichment at higher [eIF3] is expected because low-affinity RNAs consume more of the sequencing lane. **e)** Overlap of 5′-UTRs bound in 55, 166, and 500 nM libraries (red, green, and purple, respectively). See also Figure S1.

### Yeast eIF3 binds hundreds of natural 5′-UTRs with high affinity

Next, we examined which 5′-UTRs are bound with high specificity to eIF3. We focused initially on RNAs that were enriched in the 166 nM eIF3 library, which was selected as a representative of specific binding because this concentration of eIF3 was both intermediate and well correlated with adjacent concentrations (55 and 500 nM) (Figure 1c). Tracking the 10 most enriched 5′-UTRs from the 166 nM library across all other libraries revealed an enrichment score peak in the 55 nM library, which is consistent with high-affinity binding of these RNAs to eIF3 (Figure 1d). Reduced enrichment at higher concentrations of protein is expected as lower affinity binders take up more of the sequencing space (Lambert et al., 2014). 5′-UTRs were defined as “bound” or “not bound” at each concentration of eIF3 using a standard deviation-like cutoff, enrichment > (1 +range 33^rd^ to 66^th^ percentile), as previously described (Taliaferro et al., 2016). 5′-UTRs that bound non-specifically to nitrocellulose in the absence of eIF3 were eliminated from consideration (Figure S1d). Overall, we identified 1164 5′-UTRs bound in two or more adjacent concentrations of eIF3, with most of these (74%) showing binding at all three intermediate concentrations (55, 166 and 500 nM) (Figure 1e, Supplemental Table 1).

Representative 5′-UTRs, which included eIF3 binders and non-binders, were tested by nitrocellulose filter binding with purified eIF3 at concentrations ranging from 15nM to 1μM to validate the results from RBNS and determine the K_d_ of binding (Methods). Out of eleven tested binders, nine bound to eIF3 with K_d_ values from 100 to 250 nM and two did not bind tightly (K_d_ > 1000 nM) (Figures 2 and S2a-g). In parallel, we tested five non-binding 5′-UTRs (enrichment < 1.59 in 166 nM library) from the RBNS assay, all of which bound weakly or not at all with affinities above the limit for reliable quantification (K_d_ > 1000 nM) (Figure S2h-l). These results validate specific high-affinity binding of individual targets identified from comprehensive testing of eIF3 binding to yeast 5′-UTRs and show that there are hundreds of different mRNAs whose 5′-UTRs confer preferential eIF3 binding.

**Figure 2:**
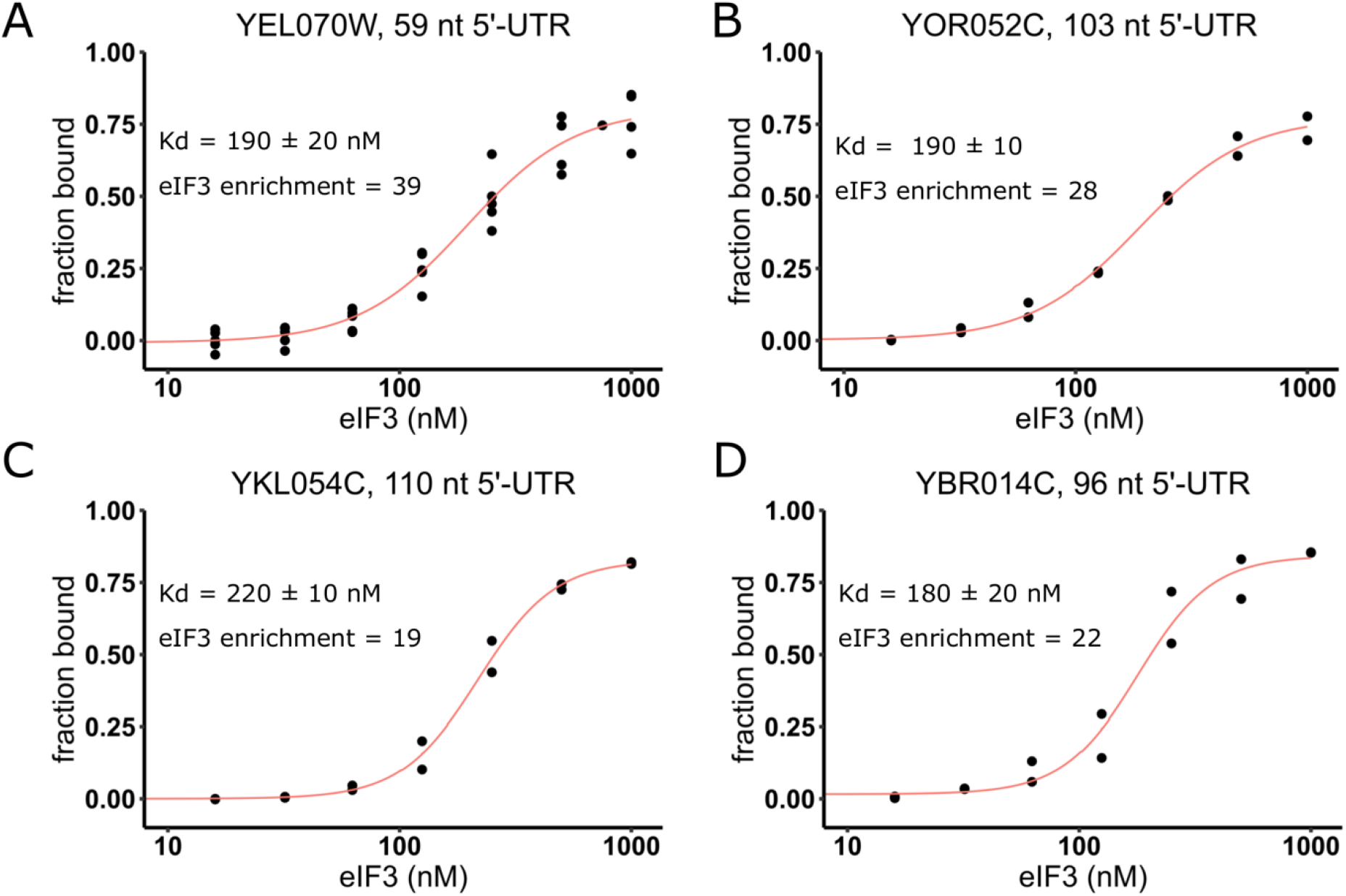
eIF3 binds to specific 5′-UTRs with nanomolar affinity. Purified eIF3 (15 – 1000 nM) was incubated with individual ^32^P-labeled 5′-UTRs from *YEL070W* **(a)**, *YOR052C* **(b)**, *YKL054C* **(c)** and *YBR014C* **(d)**. Experimental data points (black dots) were fit (red lines) to determine dissociation constants (Kd). Error reflects standard error of the fit of non-linear regression using Hill’s equation. Enrichment score at 166 nM eIF3 is shown for comparison. See also Figure S2.

### Bound 5′-UTRs are enriched in AMAYAA motifs in unstructured regions

Yeast eIF3 is composed of five distinct subunits, several of which contain known or potential RNA-binding domains, including two RNA-recognition motifs (RRM) and two helix–loop–helix (HLH) domains (Valášek et al., 2017), which have been found to mediate recognition of specific RNA sequence motifs by eIF3 and other proteins (Schuetz et al., 2014; Sun et al., 2013). Additionally, certain viral internal ribosomal entry sites (IRES) bind mammalian eIF3 with high affinity by forming an intricate RNA structure (Filbin ME et al., 2009; Walker et al., 2020). To investigate the mechanisms underlying specific binding of yeast eIF3 to a subset of 5′-UTRs, we first searched the bound RNAs from the 55, 166 and 500 nM libraries for short sequence motifs using DREME (Bailey et al., 2015). All three libraries yielded AMAYAA (where M=A or C and Y=C or U) as the most significantly enriched eIF3 binding motif with additional A-rich sequences identified in each (Figures 3a and S3a). Therefore, we focused further analysis on the AMAYAA sequence.

**Figure 3:**
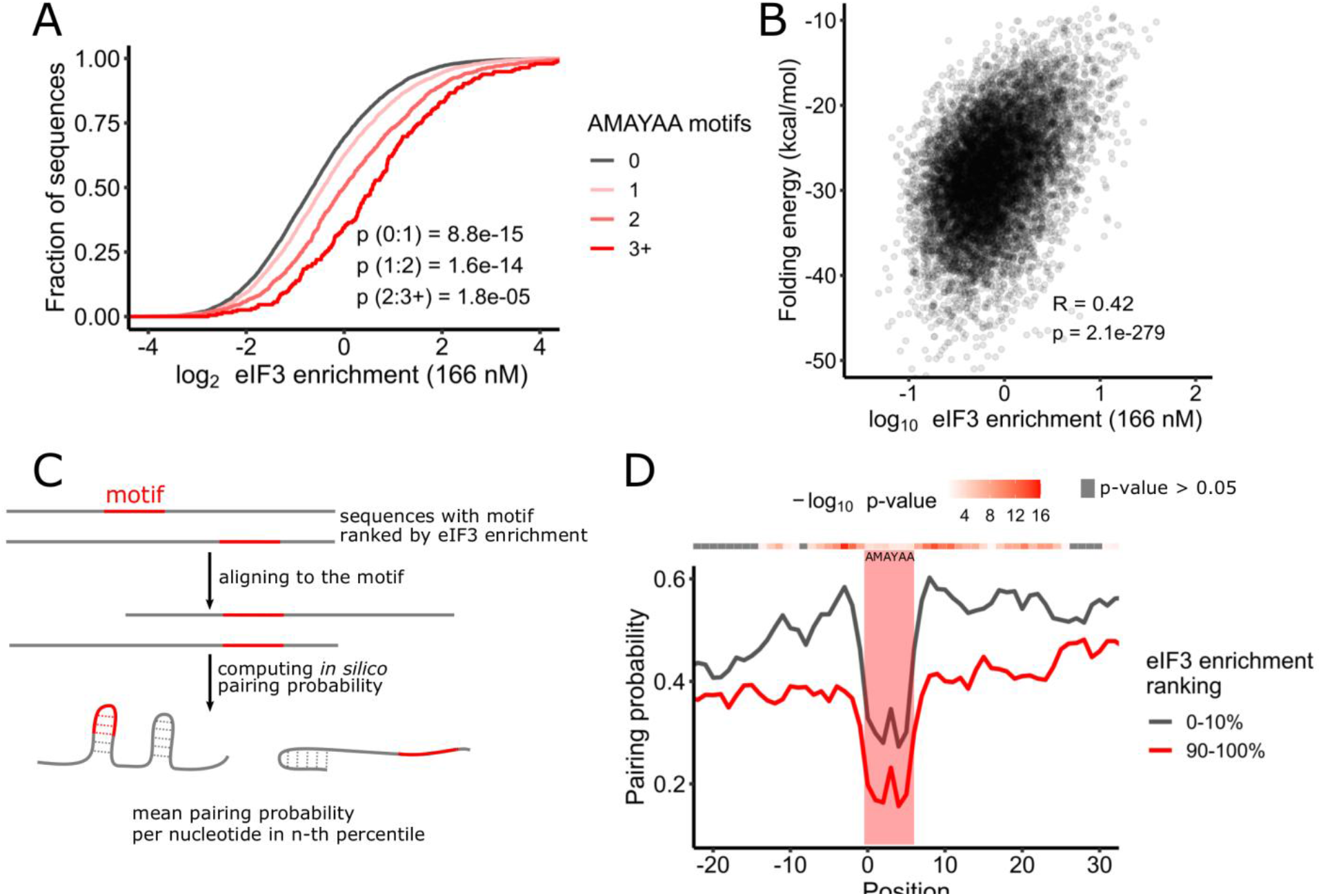
eIF3 recognizes AMAYAA motifs in unstructured regions. **a)** eIF3 binding increases with increasing numbers of AMAYAA motifs. Distribution of observed enrichment (166 nM eIF3) for 5′-UTRs with 0, 1, 2, and 3 or more AMAYAA motifs, p-adjusted (Mann-Whitney) for selected pairwise comparisons. **b)** eIF3 preferentially binds unfolded 5′-UTRs. Enrichment correlates (Pearson) with 5′-UTR folding energies calculated in RNAfold. **c)** Overview of RNA motif structure analysis. **d)** AMAYAA motifs in tight binding 5′-UTRs (top decile, red line) are more likely to be unpaired compared to motifs in weak binders (bottom decile, grey line). Average nucleotide pairing probability of 5′-UTRs binned based on their enrichment ranking in the 166 nM library. Position of the motif is indicated by the red rectangle. See also Figure S3.

If the AMAYAA motif influences eIF3 binding, we reasoned that mRNAs containing more copies of this motif would display higher affinities for eIF3. We counted the motifs in each RNA and divided the pool based on the number of motifs in the sequence. We observed that the presence of the AMAYAA motif in the RNA significantly increases enrichment score in a dose-dependent manner for sequences with 0, 1, 2, or 3+ motifs (p<1.5e-5, Figure 3b). This behavior is similar to other RNA binding proteins with known binding motifs which were analyzed using RBNS (Taliaferro et al., 2016), suggesting a bona fide sequence preference of eIF3 for AMAYAA. Overall, the enriched sequence motif AMAYAA can explain much of the observed binding specificity of purified eIF3, being present in 50.6% of binders (763 in 1508 binders).

Not all 5′-UTRs with AMAYAA motifs bind eIF3 strongly, possibly due to an additional context requirement for recognition by eIF3. Binding of yeast eIF4G to its preferred sequence motif, oligo uridine, is favored when the motif is in an unstructured context (Zinshteyn et al., 2017). We therefore hypothesized that there is a structural difference between RNAs with AMAYAA that readily bind eIF3 and those that do not. Globally, eIF3 enrichment was positively correlated with folding energy of the RNA (R=0.42, p=2.08e-279, Figure 3b), which is consistent with preferred binding of eIF3 to unstructured 5′-UTR sequences. To investigate the impact of RNA folding with nucleotide resolution, sequences containing at least one AMAYAA motif were ranked according to their enrichment score in the 166 nM library, aligned by the motif, and folded in silico using RNAfold (Figure 3c). Pairing probabilities for each nucleotide in a given bin were then averaged and plotted against nucleotide position relative to the motif. The observed nucleotide pairing probability of RNA tends to decrease with increasing enrichment (Figure S3b), with pairing probability over the motif being about 1.5-fold lower for the top decile compared to the bottom decile (Figure 3d). AMAYAA motifs are generally less likely to be paired than flanking sequences, likely due to the lack of G residues. Preferentially bound 5′-UTRs were also less folded immediately upstream and downstream of the motif. Together, these data are consistent with a preference for yeast eIF3 to bind AMAYAA motifs within unstructured regions of 5′-UTRs.

### eIF3 binders are preferentially translated in growing yeast and maintain higher translation upon glucose starvation

Ribosome footprint profiling and RNA sequencing have identified several 5′-UTR properties that correlate with translation activity genome-wide (Weinberg et al., 2016), but most of the observed variance in ribosome density still cannot be explained by current models. Thus, we asked whether direct eIF3 binding has an impact on translation in cells. Ribosome density, the average number of ribosome-protected footprints normalized to total mRNA levels, is a measure of translation activity that is thought to be predominantly affected by mRNA-specific differences in translation initiation (Shah et al., 2013). We therefore compared ribosome density in exponentially growing yeast to eIF3 binding as approximated by enrichment in the 166 nM library. We restricted this analysis to 2469 genes with a dominant 5′-UTR isoform (Methods) because ribosome footprints observed within CDS regions cannot be assigned to a specific 5′-UTR isoform and many alternative 5′-UTR isoforms show differential binding to eIF3. In fact, 913 tested genes have isoforms with significantly different (p_adj_ < 0.05) eIF3 enrichment (Figure S4a, Supplemental Table 2).

The mRNAs with 5′-UTRs that bound eIF3 in vitro (Figure 1e) showed higher ribosome density genome-wide (Figure 4a and S4b). Globally, eIF3 enrichment was positively correlated with ribosome density (R = 0.18, p=1.65e-18) (Figure 4b and S4c. In addition, mRNAs containing a greater number of AMAYAA motifs display greater ribosome density per mRNA: one motif in the 5′-UTR increased median ribosome density by 20% and two motifs increased it by 49% (Figure 4c). Together, these results are consistent with a positive influence of eIF3 binding on translation initiation for many genes.

**Figure 4:**
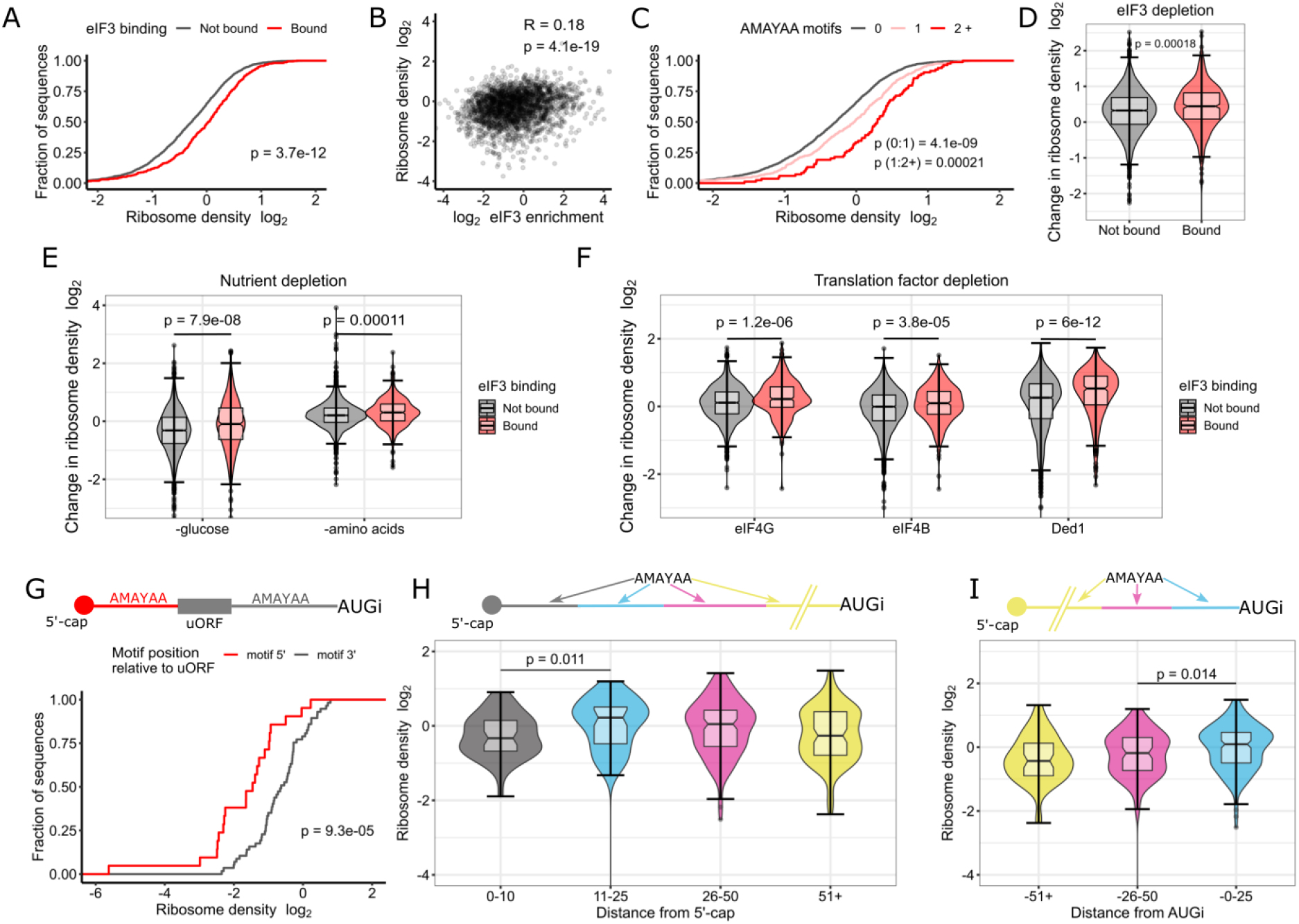
eIF3 binding and AMAYAA motifs enhance ribosome recruitment and start codon recognition in vivo. **a)** mRNAs with 5′-UTRs that bound eIF3 in vitro show higher ribosome densities in growing yeast. Cumulative distributions of ribosome densities for bound (red) or not bound (grey) mRNAs. “Bound” includes the union of 55, 166, and 500 nM libraries. Ribosome density equals the average number of ribosome-protected mRNA fragments per mRNA for genes expressing a single dominant 5′-UTR isoform (see Methods). See Table II for ribosome profiling data sources. **b)** eIF3 binding in vitro correlates with ribosome density in growing yeast (Pearson). **c)** 5′-UTR AMAYAA motifs increase ribosome density in a dose-dependent manner. **d-f)** eIF3 binding mRNAs (red) maintain higher ribosome densities than non-binding (grey) under conditions of limiting eIF3 **(d)**, following acute starvation for glucose or amino acids **(e)**, and upon inactivation of various initiation factors **(f)**. **g)** mRNAs with a 5′-UTR AMAYAA motif 5′ to a uORF (red) show reduced ribosome density compared to genes with the motif 3′ (grey). mRNAs were grouped based on the position of the first AMAYAA motif relative to the last uORF. **h,i)** Ribosome density varies by AMAYAA motif location for mRNAs without uORFs. Higher ribosome densities for mRNAs with motifs located 11-25 nt from the 5′ end of the mRNA **(h)** or 0-25 nt upstream of the start codon **(i).** Bonferroni corrected Mann-Whitney p-values shown for selected comparisons. **a-i)** all values of ribosome densities and changes in ribosome densities have been log_2_-transformed. See also Figure S4.

Next, we asked whether mRNAs with 5′-UTRs that preferentially bound eIF3 in vitro are able to maintain translation under conditions where eIF3 is limiting. We performed ribosome profiling in *tif32-td prt1-td* cells in which eIF3 levels were substantially depleted and bulk translation was reduced by ~80% (Jivotovskaya et al., 2006) (Figure S4d). The mRNAs that we observed to bind eIF3 in our RBNS experiments display greater ribosome occupancy upon eIF3 complex depletion, which is consistent with the hypothesis that they compete for limiting eIF3 under these conditions (Figure 4d). Interestingly, the apparent translational advantage of eIF3-binding mRNAs persisted in cells expressing a mutant form of eIF3i, *tif34DDKK*, which disrupts the eIF3i:eIF3b binding interface resulting in the loss of both eIF3i and eIF3g from the eIF3 complex (Herrmannová et al., 2012) (Figure S4e). This result suggests the possibility that regulation of individual eIF3 proteins could mediate mRNA-specific translational control.

We then examined translation activity of eIF3-binding mRNAs under various stress conditions where alternate mechanisms of ribosome recruitment may contribute to selective translation of some genes. We found that mRNAs which are capable of direct eIF3 binding are significantly more resistant to downregulation of translation during acute glucose starvation (p=3.7e-8) (Figure 4e) (Zid and O’Shea, 2014) and amino acid withdrawal (Figure 4e) (Santos et al., 2019). We speculate that stresses that downregulate early steps in initiation (e.g. cap-binding activity) allow selective, eIF3-dependent translation of specific mRNAs. Consistent with this possibility, eIF3-binding mRNAs maintained higher translation in cells genetically depleted of eIF4G (p=8.94e-6) as well as in cold-sensitive mutants of factors that collaborate with eIF4G to recruit ribosomes, including Ded1 (*ded1-cs*, p=8.9e-13) and eIF4B (*tif3-cs*, p=8.8e-7) (Figure 4f) (Sen et al., 2015, 2016; Zinshteyn et al., 2017). These results show that eIF3-binding mRNAs display distinct patterns of translational control.

### eIF3 binding motifs can promote or repress translation depending on their location

Our analysis of eIF3 binding in vitro and ribosome profiling in vivo is consistent with widespread translational enhancement via eIF3 binding to 5′-UTRs. However, for technical reasons, the synthetic pool used for eIF3 RBNS was limited to 5′-UTRs 122 nt. Focusing on the AMAYAA binding motif, we expanded our analysis to include all genes with a single dominant 5′-UTR and sufficient reads to quantify ribosome density in exponentially growing yeast. This allowed us to include an additional 536 genes, many of which contained multiple upstream open reading frames (uORFs). Intriguingly, while the AMAYAA motif was associated with enhanced translation if the longer 5′-UTR lacked uORFs, in mRNAs with multiple uORFs AMAYAA motifs were associated with translational repression of the main ORF (Figure S4g).

We hypothesized that upstream eIF3 binding motifs enhance translation of uORFs which leads to fewer ribosomes initiating translation of the main ORF. Additionally, as ribosomes sense the 5′-UTR sequence by scanning, we hypothesized that the order of the motif and the uORF on the 5′-UTR will matter. Therefore, we examined ribosome density on 78 mRNAs that contain one uORF and one AMAYAA motif within the 5′-UTR. Genes were separated into two categories based on the relative positions of uORF and motif – motif 5′ of the uORF or motif 3′. Notably, there was a 2-fold difference in ribosome density between mRNAs with the motif upstream vs. downstream of uORFs, with median ribosome densities of 0.24 and 0.55, respectively (p=2.7e-8) (Figure 4g). Therefore, it is likely that binding of eIF3 to AMAYAA motifs promotes recognition of downstream AUG codons, which can be in uORFs or the main ORF.

eIF3 binding to AMAYAA motifs has the potential to enhance translation initiation by multiple mechanisms which include initial recruitment of 43S pre-initiation complexes as well as recognition of initiation codons (AUG_i_) during scanning. The most likely mechanism depends on the position of the AMAYAA motif relative to the cap and AUG_i_. We therefore compared ribosome density among groups of genes with AMAYAA motifs at varying distances from the annotated transcriptional start site (TSS) and from the AUG_i_, excluding genes with uAUGs. Ribosome density was highest for mRNAs where the motif is within 11-25 nt from the TSS (Figure 4h), or within 25 nt upstream of the AUG_i_ (Figure 4i). Together, our results indicate that high-affinity binding interactions between yeast eIF3 and specific 5′-UTR sequences increase translation initiation at downstream start codons, and may do so by promoting distinct steps depending on the site of eIF3 binding (Figure 5).

**Figure 5:**
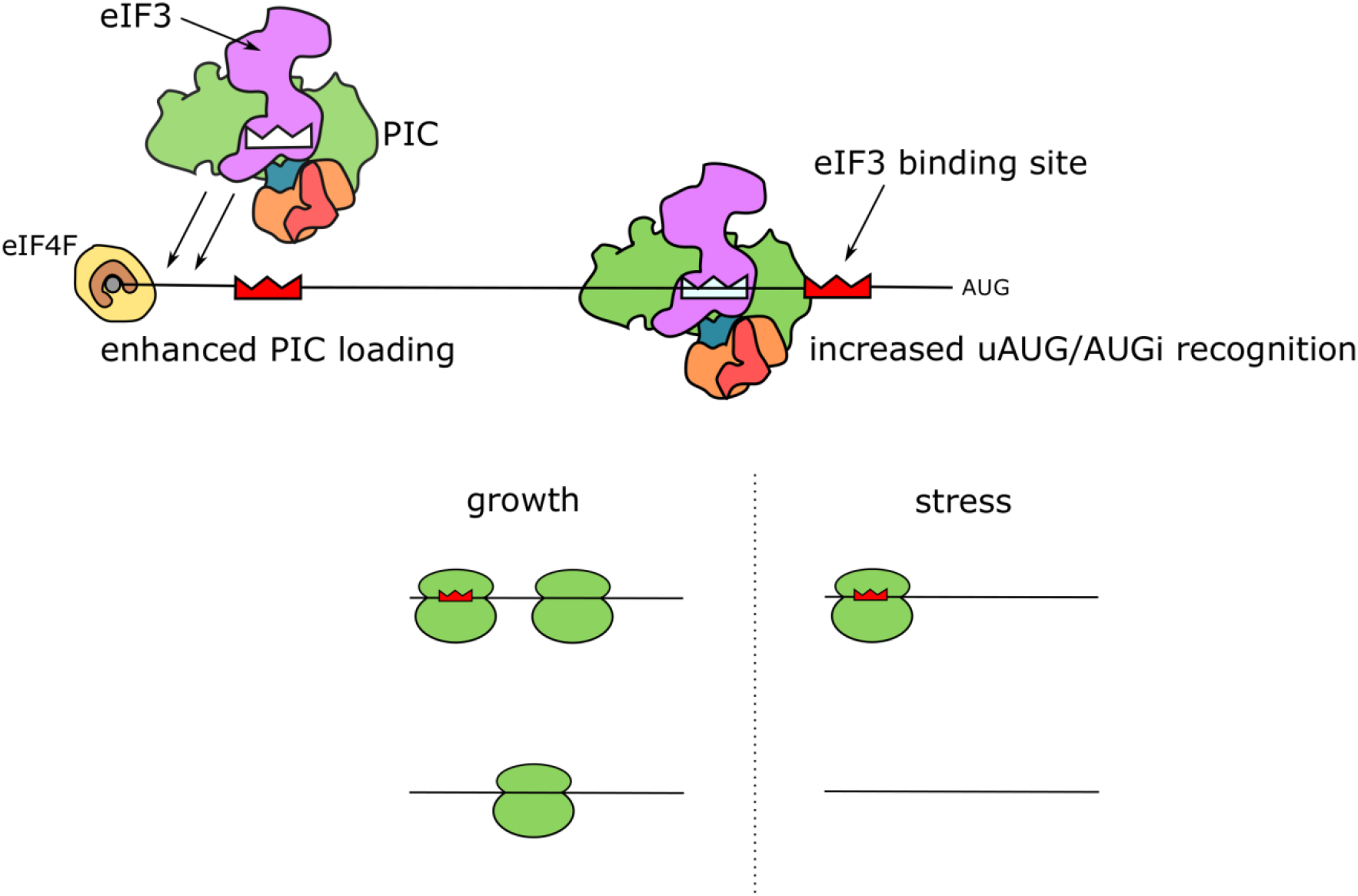
Model of location-dependent translational enhancement by direct binding of eIF3 to 5′-UTR motifs.

## Discussion

5′-UTRs are the site of action during translation initiation and are an important mRNA feature for controlling protein production post-transcriptionally in eukaryotic cells. We hypothesize that translation-enhancing elements within cellular 5′-UTRs include sequences that bind preferentially to multiple eukaryotic initiation factors (eIFs), as recently demonstrated for yeast eIF4G1 (Zinshteyn et al., 2017). Here we performed an unbiased testing of purified eIF3 binding to a library of thousands of yeast 5′-UTRs to uncover hundreds of mRNAs that preferentially bind eIF3. We show that eIF3-binding mRNAs have higher translation activity in growing cells and are less sensitive to translational inhibition in response to a variety of genetic and environmental perturbations that result in widespread effects on cellular translation. Together, our findings support a broad role for high-affinity interactions between cellular 5′-UTRs and core initiation factors for translational control of gene expression.

eIF3 promotes translation initiation by multiple mechanisms which include selective ribosome recruitment to viral mRNAs that bind eIF3 with high affinity (Filbin ME et al., 2009; Valášek et al., 2017). Our results suggest a similar role for eIF3 interactions in cellular mRNA selectivity—preferential translation of certain mRNAs under conditions that limit global initiation activity or favor non-canonical pathways of ribosome recruitment. In support, yeast eIF3-binding mRNAs maintained higher ribosome densities (average number of ribosomes per mRNA) when eIF3 protein was limited by depletion in the eIF3a/b degron strain, when cap-dependent translation was inhibited by depletion of eIF4G, and when Ded1 or eIF4B were inactivated by cold-sensitive mutations (Figure 4d, f). Globally, eIF3 binding to 5′-UTRs in vitro was modestly but significantly correlated with ribosome density in rapidly growing cells (Figure 4b and S4c). This result was surprising and suggests that binding to eIF4F is not the limiting factor for initiation on all mRNAs despite the fact that eIF4F is the least abundant initiation factor during exponential growth in rich media (Von der Haar and McCarthy, 2002). Previous work established a requirement for eIF3 for RPL41A mRNA association with PICs in vivo (Jivotovskaya et al., 2006). Our findings suggest that quantitative differences in eIF3 binding partially explain differences in PIC recruitment to different mRNAs in cells.

Analysis of eIF3-binding RNAs supports the existence of at least two modes of high-affinity binding to cellular 5′-UTRs, one that is sequence-dependent and another that remains to be determined. Binding to eIF3 in vitro (Figure 3d) and ribosome density in vivo (Figure 4c) increased with increasing numbers of AMAYAA motifs within the 5′-UTR unless the motif was located upstream of a uAUG (Figure 4g). This context-dependent effect of the eIF3-binding motif on translation of the main ORF is consistent with a simple model whereby binding to eIF3 promotes initiation on the closest downstream AUG. The observed optimal spacing with respect to the 5′ end of the mRNA is consistent with enhanced 43S recruitment immediately downstream of the cap-proximal region bound by eIF4F (Figure 4h). In addition, translational enhancement by AMAYAA motifs close to and upstream of AUGi suggests favorable interactions with 43S-bound eIF3 during start codon recognition (Figure 4i). This spacing is consistent with crosslinks observed between conserved subunits of mammalian eIF3, a and b, and the mRNA at positions - 17 to −14 relative to the start codon (Pisarev et al., 2008).

Our results raise the question of which parts of the multi-protein eIF3 complex are responsible for high-affinity binding to specific cellular 5′-UTRs. It is likely that distinct eIF3 surfaces contribute to binding in different cellular 5′-UTRs as shown for two classes of eIF3-binding viral 5′-UTRs (Neupane et al., 2020). Multiple conserved subunits of human eIF3, including a, b, d and g, crosslink directly to cellular mRNA in cultured human cells (Lee et al., 2015), highlighting the potential for distinct subunit binding preferences to contribute to mRNA selection. Candidates to mediate recognition of the AMAYAA sequence include the helix–loop– helix (HLH) motifs in eIF3a and eIF3c and the RNA recognition motif (RRM) in eIF3b. eIF3g, which also contains an RRM, appears to be dispensable for the preferential translation of many eIF3-binding mRNAs in vivo based on the observation that eIF3-binding mRNAs maintain high ribosome density in *tif34DDKK* mutants (Figure S4f) in which eIF3g and eIF3i are destabilized from the core complex of eIF3a/b/c (Herrmannová et al., 2012). Our results also suggest the potential for mRNA-specific translational regulation by post-translational modifications to specific domains within eIF3, such as succinylation of the RRM in eIF3b or phosphorylation of the RRM in eIF3g (Albuquerque et al., 2008; Weinert et al., 2013).

The comprehensive approach used here to identify cellular 5′-UTRs that bind to wild-type yeast eIF3 can be used to tease out the contributions of individual domains, and even specific amino acids of interest. It is increasingly clear that core initiation factors engage cellular 5′-UTRs in highly specific interactions that contribute to mRNA-specific rates of initiation—in growing cells and global re-tuning of translation during stress. Our results suggest a broad potential for direct eIF3 binding to maintain translation of specific cellular mRNAs during a variety of stress responses. Such mRNA-specific sensitivities are likely to be important for the pathological effects of dysregulated initiation factors in cancer and other human diseases.

## Acknowledgements

We thank Alan Hinnebusch and Jon Lorsch for the eIF3 production strain and eIF3 purification protocols. We thank members of the Gilbert lab for discussion and comments on the manuscript. This work was supported by NIH R01GM132358 to W. Gilbert and R15GM140372 to C. Aitken. R. Niederer was supported by the American Cancer Society (Postdoctoral Fellow) and the NIH (K99GM135533).

## Supplemental Figures

**Figure S1:**
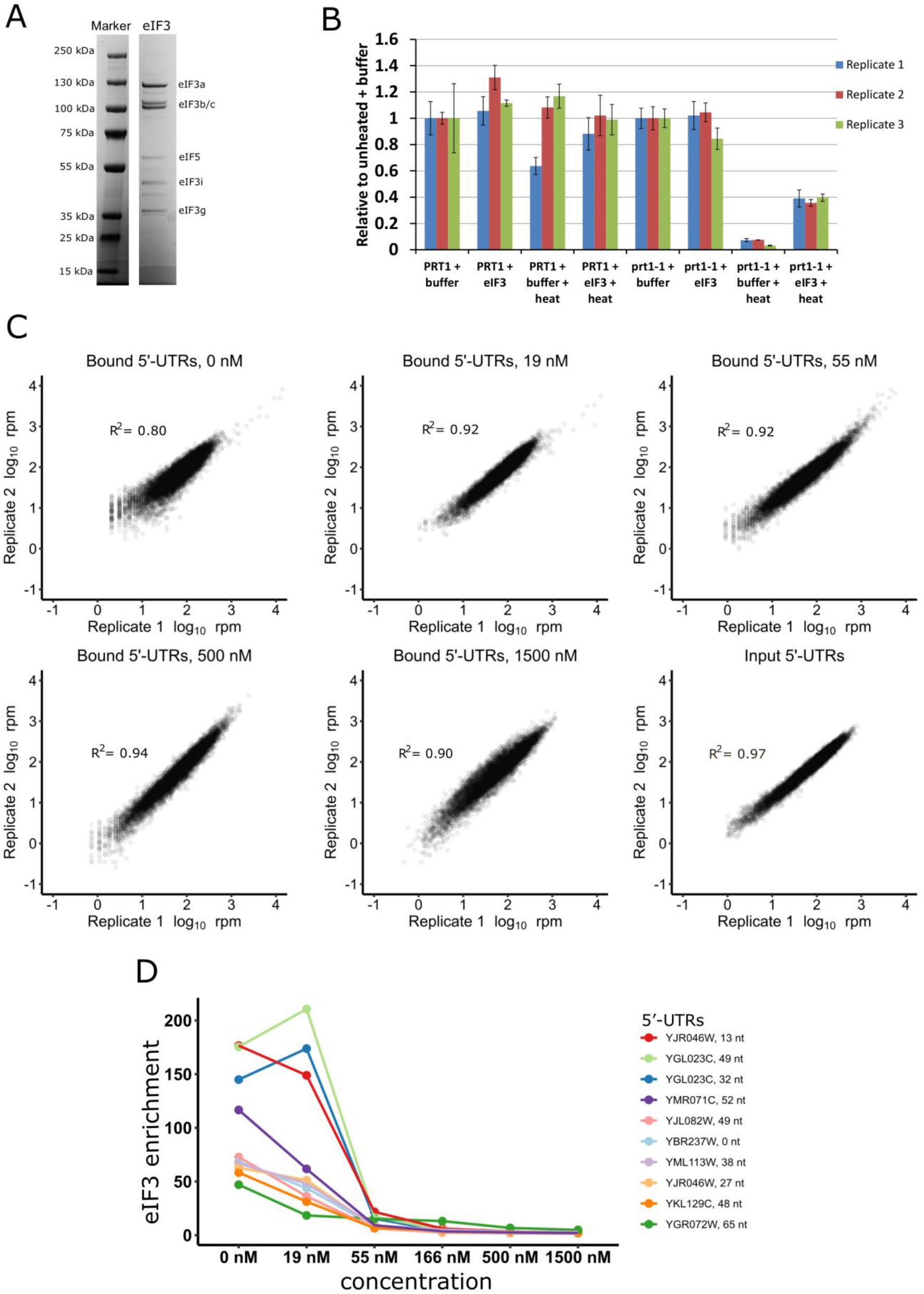
Control experiments for eIF3 RNA Bind-n-Seq. **a)** Commassie blue-stained tris-glycine gel of purified eIF3 with indicated subunits. The major contaminant observed is eIF5. **b)** Purified eIF3 restores translational activity to heat-inactivated extracts from *prt1-1* mutant yeast. Reported are the relative luciferase activities of eIF3 heat sensitive (*prt1-1*) and isogenic (*PRT1*) yeast extracts supplemented with either 100 nM purified eIF3 in storage buffer (+eIF3) or equal volume of storage buffer (+Buffer). Replicates indicate independently prepared extracts. Bars represent the average of 3 technical replicates and error bars represent standard deviation. **c)** RBNS is reproducible for all libraries. Shown are the bound reads for libraries at 0, 19, 55, 500, and 1500 nM eIF3, as well as the reads for the input libraries. **d)** Enrichment of top 10 binders in 0 nM libraries traced across all libraries. The gradual decrease in eIF3 enrichment score with increased eIF3 concentration is consistent with constant background binding with increasing amount of specifically bound RNA (Lambert et al., 2014).

**Figure S2:**
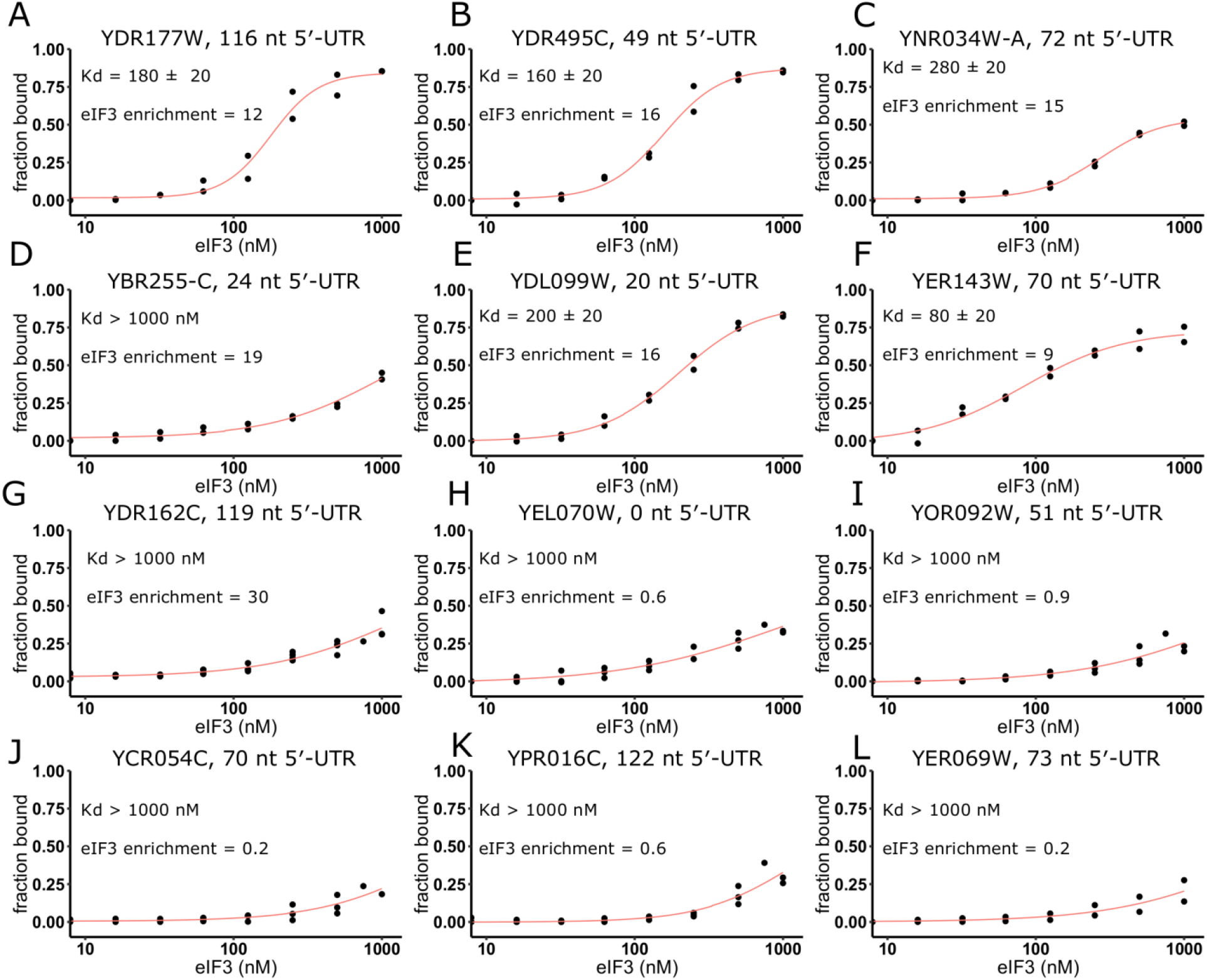
eIF3 enrichment score is predictive of eIF3 binding affinity. Increasing amounts of eIF3 were bound with ^32^P-labeled 5′-UTRs from **(a)** *YDR177W*, **(b)** *YDR495C*, **(c)** *YNR034W−A*, **(d)** *YBR255C−A*, **(e)** *YDL099W*, **(f)** *YER143W*, **(g)** *YDR162C*, **(h)***YEL070W*, **(i)** *YOR092W*, **(j)** *YCR054C*, **(k)** *YPR016C* or **(l)** *YER069W*. 5′-UTRs in **(a-g)** represent binders in the RBNS assay (eIF3 enrichment > 1.59) and **(h-l)** represent non-binders. Apparent dissociation constant calculated from non-linear regression using Hill’s equation in Origin is indicated together with the obtained eIF3 enrichment score.

**Figure S3:**
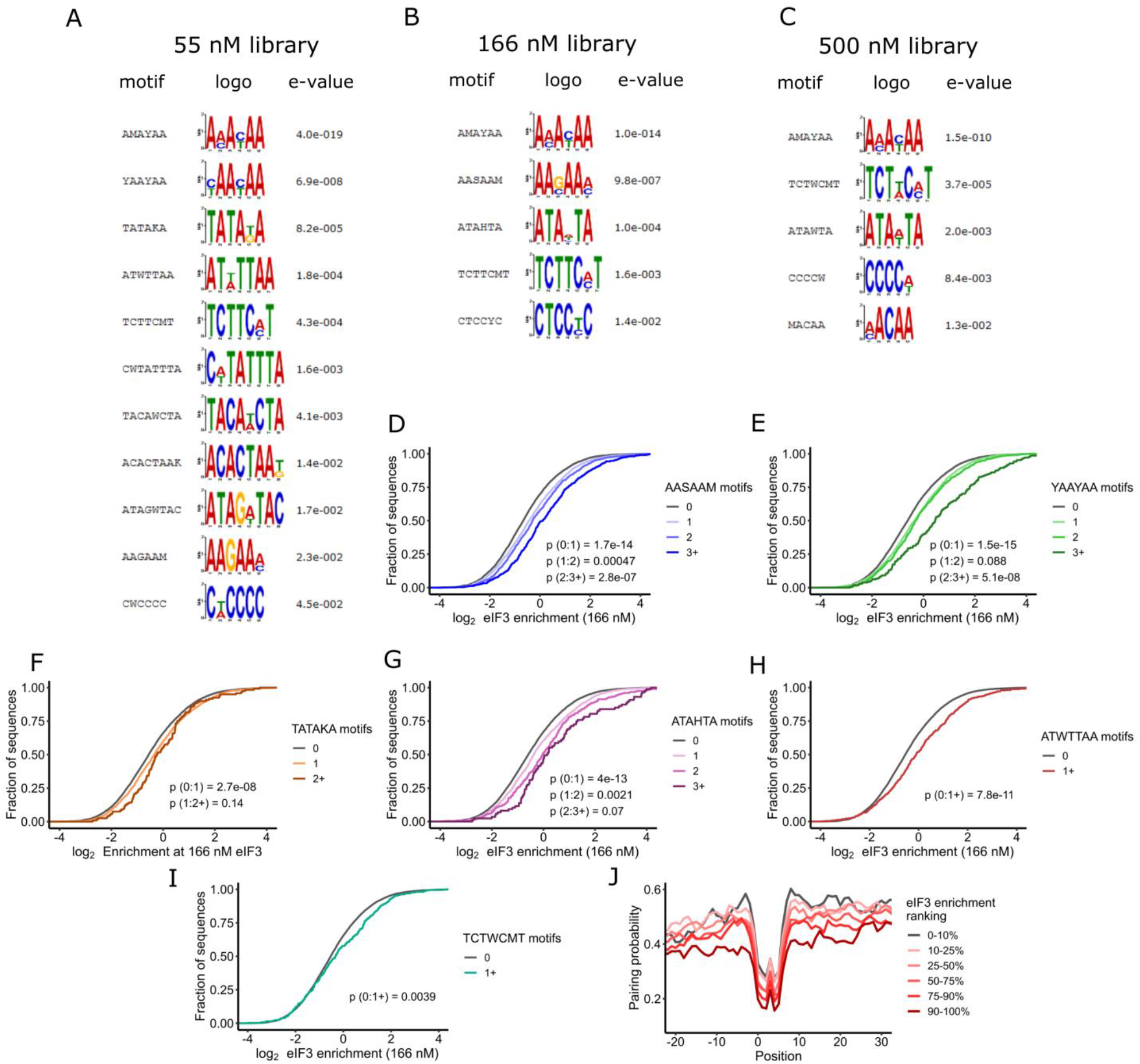
eIF3 preferentially recognizes AMAYAA motifs in unstructured regions. Sequence motifs identified by DREME (Bailey et al., 2015) in **(a)** 55 nM, **(b)** 166 nM or **(c)** 500 nM as overrepresented among bound 5′-UTRs. E-value represent Fisher’s exact test p-value for a given motif multiplied by number of motifs tested. Only motifs with e-value < 0.05 are shown. Distribution of observed enrichment in 166 nM library is shown for motifs **(d)** ASSAAM, **(e)** YAAYAA, **(f)** TATAKA, **(g)** ATAHTA, **(h)** ATWTTAA, or **(i)** TCTWCMT. **(j)** Higher eIF3 enrichment scores for 5′-UTRs containing AMAYAA motif are associated with lower RNA base-pairing probabilities of the motif and neighboring nucleotides.

**Figure S4:**
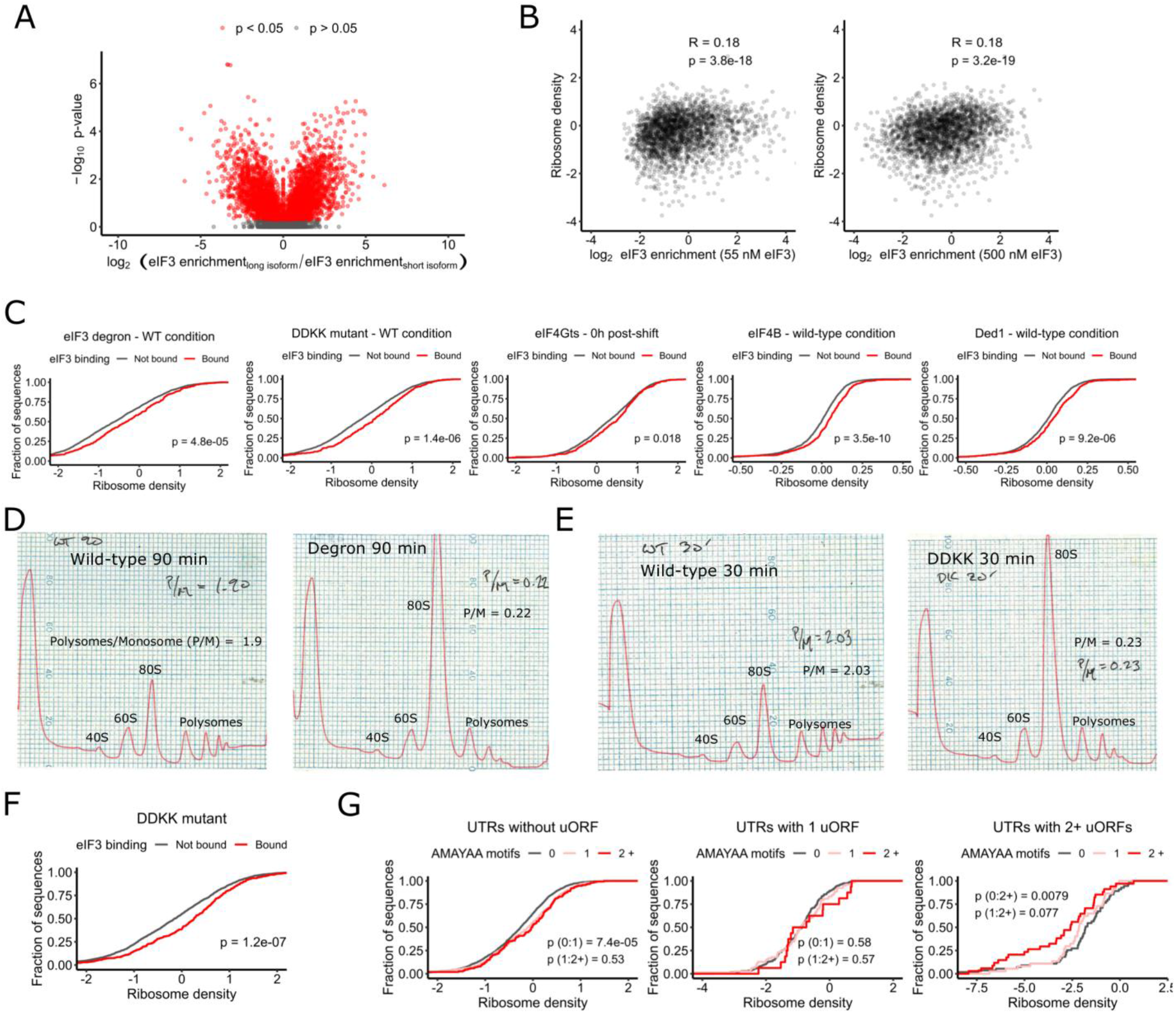
eIF3 binding and AMAYAA motifs enhance ribosome recruitment in vivo. **a)** 5′-UTR isoforms from hundreds of genes show differential binding to eIF3. Differential eIF3 enrichment for short and long isoform is plotted together with Bonferroni corrected p-values (t-test). Significant differences p_adj_ < 0.05 are depicted in red. **b)** eIF3 enrichment in 55 and 500 nM libraries is correlated with ribosome densities. eIF3 binding is associated with increased ribosomal densities across multiple datasets in **(c)** wild type or pseudo-wild type and in **(f)** eIF3 DDKK mutant cells. eIF3 depletion leads to > 80 % decrease in translation in **(d)** eIF3 degron or **(e)** DDKK mutant cells as estimated from the change of polysome/monosome (P/M) ratio. **(g)** The AMAYAA motif is associated with increased ribosome density in 5′-UTRs without uORFs. In 5′-UTRs with multiple uORFs, the AMAYAA motif is associated with decreased ribosome density. **b, c, f, g)** Depicted are the log_2_-transformed values of ribosome densities.

## Material and methods

### Yeast strains and growth

Genotypes are listed in Table I. Strain LPY87 was grown for eIF3 purification as previously described (Phan et al., 2001). Briefly, culture was grown overnight in synthetic complete (SC) media without leucine and uracil at 30°C. The starter culture was used to inoculate 18 L of YPD media and grown for 14 – 18 hours to OD_600_ 4 – 5. Wild type (*PRT1*) and eIF3 temperature-sensitive (*prt1-1*) strains used to prepare extracts for in vitro translation and complementation assays were grown at 23 °C in YPD. The *eIF3a/b* degron strain YAJ34 (Jivotovskaya et al., 2006) was grown at 25 °C in SC_Raff_ + Cu^2+^ before being shifted to pre-warmed SC_Raff/Gal_ + BCS at 36 °C for 90 min to deplete eIF3a/b. The temperature-sensitive *tif34-DDKK* strain (Herrmannová et al., 2012) was grown at 30°C in SC media before being shifted to pre-warmed SC media at 37 °C for 30 min.

**Table I:**
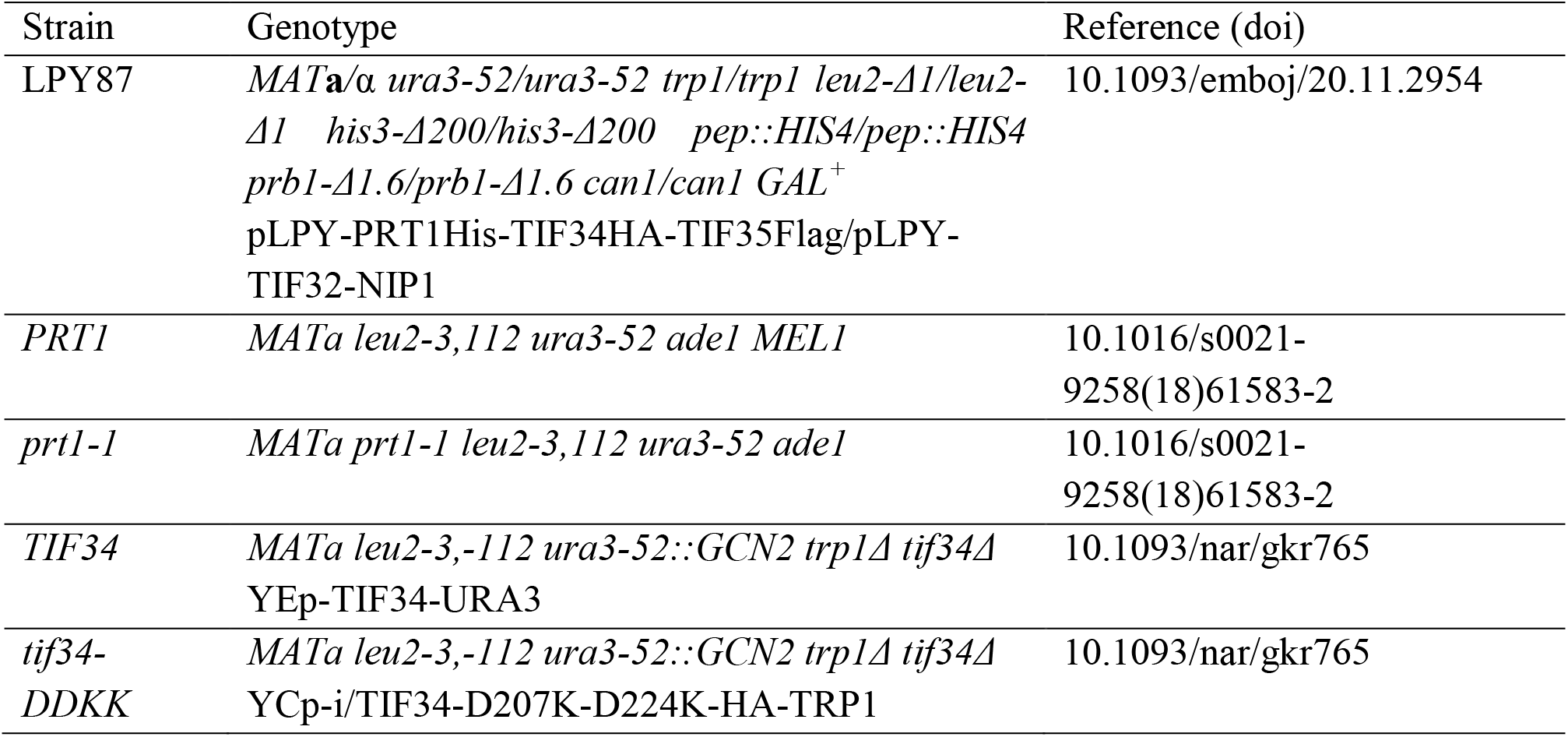
Yeast strains

**Table II:** Datasets

| Name | Authors | Description | GEO accession number | Year |
| --- | --- | --- | --- | --- |
| Promoter sequences direct cytoplasmic localization and translation of mRNAs during starvation in yeast. | Zid BM, O'Shea EK | ribosome profiling of glucose-starved yeast | GSE56622 | 2014 |
| Genome-wide analysis of translational efficiency reveals distinct but overlapping functions of yeast DEAD-box RNA helicases Ded1 and eIF4A | Sen ND, Zhou F, Ingolia NT, Hinnebusch AG | ribosome profiling of Ded1-depleted yeast | GSE66411 | 2015 |
| Improved Ribosome-Footprint and mRNA Measurements Provide Insights into Dynamics and Regulation of Yeast Translation | Weinberg DE, Shah P, Eichhorn SW, Hussmann JA, Plotkin JB, Bartel DP | ribosome profiling of wild-type yeast | GSE75897 | 2016 |
| eIF4B stimulates translation of long mRNAs with structured 5' UTRs and low closed-loop potential but weak dependence on eIF4G | Sen ND, Zhou F, Harris M, Ingolia NT, Hinnebusch AG | ribosome profiling of eIF4B-depleted yeast | GSE81966 | 2016 |
| Translation initiation factor eIF4G1 preferentially binds yeast transcript leaders containing conserved oligo-uridine motifs | Zinshteyn B, Rojas-Duran MF, Gilbert WV | ribosome profiling of eIF4G-depleted yeast | GSE87614 | 2017 |
| Cycloheximide can distort measurements of mRNA levels and translation efficiency | Santos DA, Shi L, Tu BP, Weissman JS | ribosome profiling of amino acid-starved yeast | GSE125038 | 2019 |

### eIF3 purification

LPY87 cells expressing His-tagged eIF3 were harvested (85-95 g of wet cell pellet), washed with lysis buffer (20 mM HEPES.KOH, 350 mM KCl, 5 mM MgCl2, 10 % Glycerol, 20 mM Imidazole, 10 mM Beta-mercaptoethanol, pH = 7.4), resuspended in 30 mL of lysis buffer, frozen as droplets in liquid N2 and stored in −80°C until lysis on Retch cryomill, using 2 × 5 min shaking at 15 Hz with 1 min intermittent cooling at 5 Hz. Cell powder was thawed in 400 mL of lysis buffer in the presence of cOmplete protease inhibitors (Sigma-Aldrich), 1 μg/ml pepstatin A, 1 μg/ml aprotinin, 1 μg/ml leupeptin, and 100 μL of Turbo DNase. Lysate was clarified at 12,500 × g for 40 min and applied to 5 mL of freshly regenerated Ni 2+ Sepharose. Bound protein was washed with lysis buffer until there was no detectable protein in the flow through and eluted using lysis buffer supplemented with 350 mM Imidazole. Fractions containing eIF3 were concentrated with 10 kDa MWCO centrifugation columns to ~ 2 mL and resolved in two batches on HiLoad 16/60 Superdex 200 preequilibrated in low salt buffer 20 mM HEPES.KOH, 100 mM KCl, 10 % glycerol, 0.1 mM EDTA, 2 mM DTT, pH = 7.4. Fractions containing all 5 eIF3 subunits were combined and loaded on phosphocellulose column prepared from cellulose phosphate (Sigma-Aldrich) sequentially activated in 1M HCl, 1 M NaOH and preequilibrated in low salt buffer. The column was then washed with 100 mL of low salt buffer and the protein was sequentially eluted with 200/350/1000 mM KCl in 20 mM HEPES.KOH, 10 % glycerol, 0.1 mM EDTA, 2 mM DTT, pH = 7.4. eIF3-containing fractions were pooled dialyzed against 2 L of storage buffer (20 mM HEPES.KOH, 100 mM KOAc, 10 % Glycerol, 2 mM DTT, pH = 7.4), concentrated to 0.5 – 1 mL and stored in 15 μL aliquots in −80°C. Final eIF3 concentration was determined using Bradford assay with BSA as the protein standard.

### eIF3 complementation assay

Translationally active extracts were prepared from wild type (*PRT1*) and heat sensitive eIF3 mutant (*prt1-1*) strains grown at 23 °C using published protocols (Zinshteyn et al., 2017). Three replicate extracts prepared from independent cultures were either pre-treated at 39 °C for 10 minutes or kept on ice. Translation of capped nanoluciferase reporter mRNA was performed with supplemental 100 nM eIF3 or buffer control in three technical replicates as described previously (Rojas-Duran and Gilbert, 2012). The reaction was stopped after 30 minutes by 100-fold dilution into PBS and the amount of nanoluciferase was measured using Nano-Glo (Promega) on a Centro XS3 luminometer (Berthold).

### 5′-UTR pool design and synthesis

5′-UTR boundaries and abundances were calculated from sequencing of wild type yeast (Pelechano et al. 2014). When a 5′-UTR started within 10 nts of its nearest neighbor, the sequences were merged. Inclusion in the pool also required the following: 5′-UTRs must be expressed within 25% of the mode abundance for a given 5′ UTR, and 5′-UTRs must make up at least 5% of the total abundance for that ORF, unless the mode was <5% of the total, in which case we used the mode. Upstream AUGs within 761 (6.3% of all) 5′-UTR sequences were mutated to AGT such that the first AUG encountered by a scanning pre-initiation complex moving 5′ to 3′ would be the annotated AUGi. Each sequence consisted of a randomized 10 nucleotide unique identifier barcode and an adaptor sequence used for priming reverse transcription and Illumina sequencing. RNA was *in vitro* transcribed from the PCR-amplified pool using in house prepared T7 RNA polymerase and gel-purified.

### RBNS library preparation

Binding reactions (50 μL) were assembled at room temperature with 60 ng/μL of pool RNA (~1 μM) and various concentrations of eIF3 (0-1500 nM, 2 replicates each) in 20 mM HEPES. KOH, 100 mM KOAc, 2 mM Mg(OAc)2, 10 % glycerol, 2 mM DTT, RNAseIn (80 U, Promega), and 100 ng/μL of yeast tRNA (~4 μM, Sigma-Aldrich). After 30 minutes, the reaction was passed through layered nitrocellulose (top) and nylon (bottom) filters preequilibrated in binding buffer using a vacuum manifold. Filters were washed 2 × 200 μL with ice-cold binding buffer and dried. eIF3-bound RNA was extracted from nitrocellulose by incubation with proteinase K (Sigma-Aldrich), concentrated by Zymo column, and reverse transcribed with AMV reverse transcriptase (RT primer: OWG921). Gel-purified cDNA was ligated with a barcoded (10N) 5′-adapter as described (Niederer et al., 2020).

### Ribosome profiling of eIF3a/b degron and eIF3i-DDKK

Each strain and its matching wild type (Table I) were grown under restrictive conditions (see Yeast strains and growth) for a duration resulting in ~80% decrease in bulk translation as judged by analytical polysome profiling and quantification of polysome:monosome ratios as previously described (Jivotovskaya et al., 2006) Cycloheximide was added to a final concentration of 100 μg/mL 2 min prior to harvesting by filtration through a Kontes filtration apparatus and flash freezing in liquid nitrogen. We performed all subsequent steps as described previously (Sen et al., 2016).

### Data analysis

RBNS raw sequencing reads were trimmed with cutadapt and aligned to the pool sequences. PCR duplicates in each library were removed using 10N barcodes in the 5′-adapter and reads were normalized in each library according to their sequencing depth. Artificial sequences designed for other studies were excluded from the data processing and analysis (Niederer et al., 2020). For quantification, we required >1 rpm in each of the input replicates and >2 reads in each of the sample libraries. Enrichment score for each 5′-UTR in a given library was then calculated as the ratio of normalized reads in the library to input. Data was processed and visualized in Python and R using custom scripts.

For comparisons between translation activity in vivo and eIF3 binding in vitro, in vivo 5′-UTRs were defined based on sequencing reads (Pelechano et al., 2014). For every ORF, 5′-UTRs within 10 nts of one another were merged. Only 5′-UTRs that contributed at least 5% to the total mRNA pool for a given gene were considered. Dominant 5′-UTRs were defined as the most abundant 5′-UTR for a given gene, which must account for at least 40 % of all mRNAs for that gene and be at least twice as abundant as the second most common 5′-UTR. The presence of an upstream AUG was the selection criteria for assigning uORFs to a given 5′-UTR. For the motif positional comparisons, only 5′-UTRs with sizes between 40 and 150 nucleotides were selected.

### Filter binding

Individual 5′-UTRs were amplified from the pool using target-specific primers and in vitro transcribed. Gel-purified RNA was dephosphorylated (FastAP, ThermoFisher) and 5′-labeled with 32P-γ-ATP (Perkin Elmer) using T4 Polynucleotide kinase (NEB). Binding reactions (5 μL) contained ~10 nM labeled RNA and 0/16/32/63/125/250/500/1000 nM eIF3 in buffer as described for RBNS. After 30 minutes, reactions were passed through layered nitrocellulose (top) and nylon (bottom) filters preequilibrated in binding buffer. Filters were washed with 80 μL of ice-cold binding buffer, disassembled and dried. Captured RNA was visualized by phosphorimaging and quantified using ImageJ (Schneider et al., 2012). At each concentration, eIF3-bound RNA was quantified as the ratio of nitrocellulose-bound RNA to the sum of RNA captured on both filters. Data were fit using nonlinear regression with Hill’s equation in Origin.

